# Cycle-consistent deep generative modeling unifies cellular states across unpaired spatial and single-cell modalities

**DOI:** 10.64898/2026.05.25.727736

**Authors:** Haoran Zhang, Jeffrey F. Quinn, Data Science TeamLab, Wesley Tansey

**Affiliations:** Department of Computer Science, University of Texas at Austin; Computational Oncology, Department of Epidemiology & Biostatistics, Memorial Sloan Kettering Cancer Center; Break Through Cancer

## Abstract

Current spatial and single-cell technologies capture complementary but incomplete views of cellular state, with transcriptomic, proteomic, and spatial information distributed across distinct platforms. Integration is challenged by unpaired measurements, mismatched feature spaces, and modality-specific biases. We present MultiTME, a multimodal framework that integrates heterogeneous spatial and single-cell data using a spatially-regularized, cycle-consistent deep generative model. By enforcing consistency of bidirectional mappings, MultiTME learns a shared latent representation that enables translation between modalities without requiring paired observations or shared features. Across benchmarks, MultiTME outperforms existing methods, produces accurate cross-modal cell typing, improves spatial transcriptomic panel completion, and transfers whole-transcriptome information to generate spatially resolved maps at cellular resolution. Applied to a multimodal colorectal cancer dataset, we demonstrate that MultiTME integration reveals a spatially coherent proliferative–invasive tumor axis not directly observable within single modalities. Across five multimodal spatial datasets, we show MultiTME can correct for platform-specific biases between Xenium and CosMx, thereby facilitating cross-dataset harmonization and enabling pan-cancer spatial studies. Code for MultiTME is available at https://github.com/tansey-lab/multitme.

## Introduction

The tissue microenvironment (TME) is a spatially organized system in which cellular composition, molecular state, and spatial architecture jointly determine tissue function and disease progression^1,2^. Recent advances in spatial and single-cell technologies have enabled quantitative profiling of the TME, but each modality captures only a partial view of cellular state ^3,4^. Imaging-based proteomics offers high spatial resolution, but limited panel size, imaging-based spatial transcriptomics measures targeted gene sets or whole transcriptomes in spatially resolved bins with modality-specific artifacts ^5,6,7^, and scRNA-seq provides genome-wide profiles without spatial context. As a result, each dataset represents a modality-specific projection of an underlying biological state that is not directly observed, and recovering such a state requires integrating measurements across assays.

Integration is most straightforward when modalities are measured in the same cells, but in practice, such paired measurements are rare. Most spatial and single-cell datasets are acquired on different cells, adjacent tissue sections, or in separate experiments, with no shared cellular correspondence among them ^8^. Many of the existing analytical methods were developed under assumptions of paired measurements ^9,10^ or feature overlap ^11,12^, leaving misaligned and non-overlapping modalities poorly supported. This mismatch between methodological assumptions and experimental reality has led to calls for methods that integrate modalities through a shared underlying biological state that places emphasis on modeling distributions rather than cells. ^8^

MExisting integration approaches only partially address this challenge. Recent methods have been developed that leverage reference-based mapping ^13,14,11^, pairwise matching of shared biomarkers ^15,16^, and graph-based integration ^17,18^. While effective in specific settings, these methods either rely on observations of multimodal measurements from the same cells or require expert knowledge to connect features across modalities. Further, when integrating spatial modalities, many methods treat cells independently and underutilize spatial context as a structural constraint ^1^. The key methodological gap in the field is a need for methods that can flexibly integrate unpaired modalities across both single-cell and spatial technologies.

Here we introduce MultiTME, a deep generative framework for integrating unpaired multimodal single-cell and spatial measurements. MultiTME defines modality-specific observation models coupled through a common representation and shared latent space. A novel cyclic loss function encourages the MultiTME latent space to remove technical differences between modalities and instead focuses it on shared biological states. Optional auxiliary classification and spatial losses allow MultiTME to incorporate expert cell typing and spatial context directly into learning. The resulting model produces a coherent integration of heterogeneous spatial and single-cell datasets without requiring paired observations or external supervision. Across multiple platforms, including Xenium, CosMx, Visium HD, CODEX, and scRNA-seq, MultiTME improves cell type identification, modality completion, and cross-platform translation, providing a unified framework for the integration and interpretation of the tissue microenvironment.

## Results

### MultiTME learns a common latent space across modalities via cycle consistency mechanism

Spatial and single-cell assays capture complementary molecular views of tissue but typically measure different features and are often acquired from serial sections rather than identical cells. MultiTME addresses this challenge by learning a shared latent space where cells from different modalities become directly comparable, enabling cross-modal cell type assignment, spatial analysis across profiling technologies, and prediction of unmeasured modalities. The model consists of three components: modality-specific projections, a shared latent encoder, and modality-specific decoders. Each modality is first projected into an intermediate representation in a common dimensionality through modality-specific projectors, allowing the model to accommodate differences in feature dimensionality and measurement scales across technologies (e.g. tens of proteins in spatial proteomics versus thousands of genes in transcriptomic assays). The projected representations are then mapped by a shared encoder that generates a low-dimensional latent representation *z*. The encoder contains no modality indicator, encouraging the learned representation to capture the common biological state rather than platform-specific artifacts. Observations are reconstructed through modality-specific decoders.

To translate information across modalities without requiring paired measurements, MultiTME enforces cycle consistency across modalities. Starting from a cell measured in one modality, the model first encodes it into the shared latent space and then decodes it into the feature space of another modality, producing a predicted observation in the second modality. The predicted observation is then re-encoded and mapped back to the original modality (Figure 1b). The model is trained so that this cycle returns both the latent representation and the reconstructed observation close to the original input. By enforcing consistency at both the latent and observation levels, the model learns modality-invariant biological representations while maintaining accurate modality-specific reconstructions.

**Figure 1:**
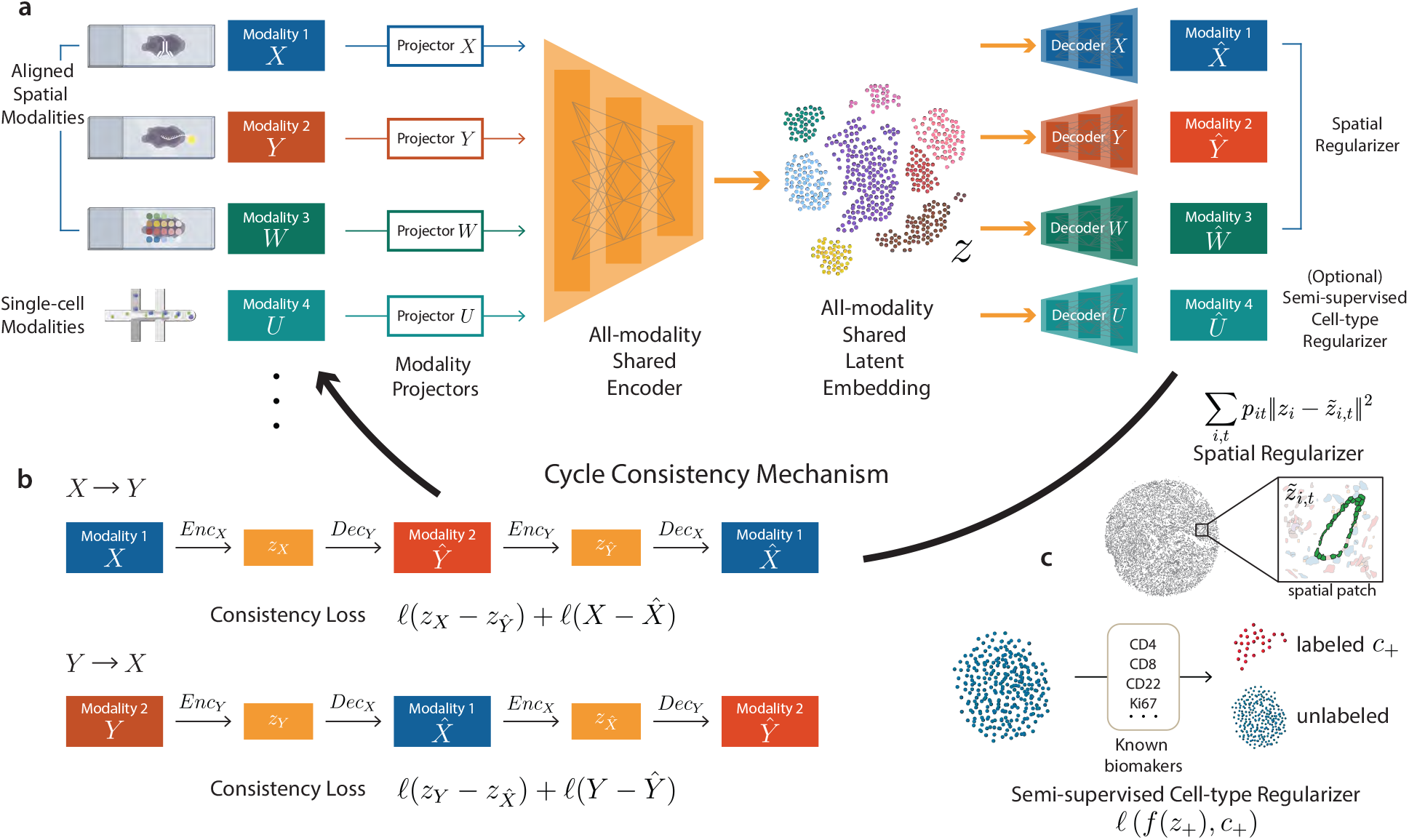
Overview of the MultiTME framework for integrating unpaired spatial and single-cell modalities. (**a**) Multiple input modalities are mapped through modality-specific projectors into a shared latent representation that captures common biological structure while accommodating modality-specific measurement differences. (**b**) The cycle consistency mechanism aligns latent representations across modalities without paired supervision. (**c**) The spatial regularizer enforces that the latent representation of a cell is aligned with the weighted mean of cells of the same type in the local neighborhood (green). The semi-supervised cell type regularizer preserves the biological interpretability of the learned latent space.

Spatial information is incorporated through a neighborhood regularization mechanism (Figure 1c). For modalities with spatial coordinates, local neighborhoods are constructed using k-nearest neighbors in tissue space (see Methods). A graph fusion penalty encourages cells of the same type that are spatially close (e.g., green cells in Figure 1c) to have similar latent representations, reflecting the biological observation that neighboring cells often share related states due to tissue structure and microenvironmental interactions. This spatial regularization stabilizes representation learning and helps preserve spatial organization in the integrated latent space.

When annotations are available in one modality, MultiTME incorporates semi-supervised cell type guidance to anchor the latent representation to biologically meaningful structure. For unlabeled modalities, the model identifies high-confidence examples of specific cell types using canonical marker genes or proteins (see Methods). These pseudo-labeled cells provide weak supervision through a classification head attached to the latent representation (Figure 1a,c). This hybrid strategy helps organize the latent space according to biologically interpretable cell identities and improves downstream tasks such as cell type annotation and modality translation.

Together, these components enable MultiTME to learn a unified representation across heterogeneous modalities without requiring paired measurements. The learned latent space captures the common underlying biological state while retaining modality-specific decoding, enabling downstream tasks including cross-modal translation, panel completion, and multimodal cell type characterization.

### MultiTME enables spatially guided cell typing across spatial modalities

We first evaluated MultiTME on the task of cell typing in spatial multi-omic data, the foundational step for interpreting tissue organization and cellular interactions. To demonstrate this capability, we analyzed a human tonsil dataset ^19^ with scRNA-seq and CODEX spatial proteomics measurements from the same patient. The scRNA-seq dataset provides whole-transcriptome profiles and cell type annotations, while the CODEX assay captures spatially resolved protein measurements across the tissue. MultiTME integrates these modalities by learning a shared representation of cellular states that leverages transcriptomic reference information to automatically annotate CODEX cells while preserving spatial constraints (Figure 2a). Visualizing the shared embedding space via a joint UMAP projection, cells from scRNA-seq and CODEX have consistent transcriptional manifolds, indicating that MultiTME successfully aligns dissociated and spatial measurements (Figure 2b). Using this shared representation, MultiTME accurately reconstructs the spatial organization of major immune populations in the tonsil. The predicted spatial cell type map closely matches the pathologist annotations in CODEX, capturing canonical lymphoid architecture including germinal-center B cells, proliferating B cells, CD4 T cells, CD8 T cells, dendritic cells, and plasma cells (Figure 2c).

**Figure 2:**
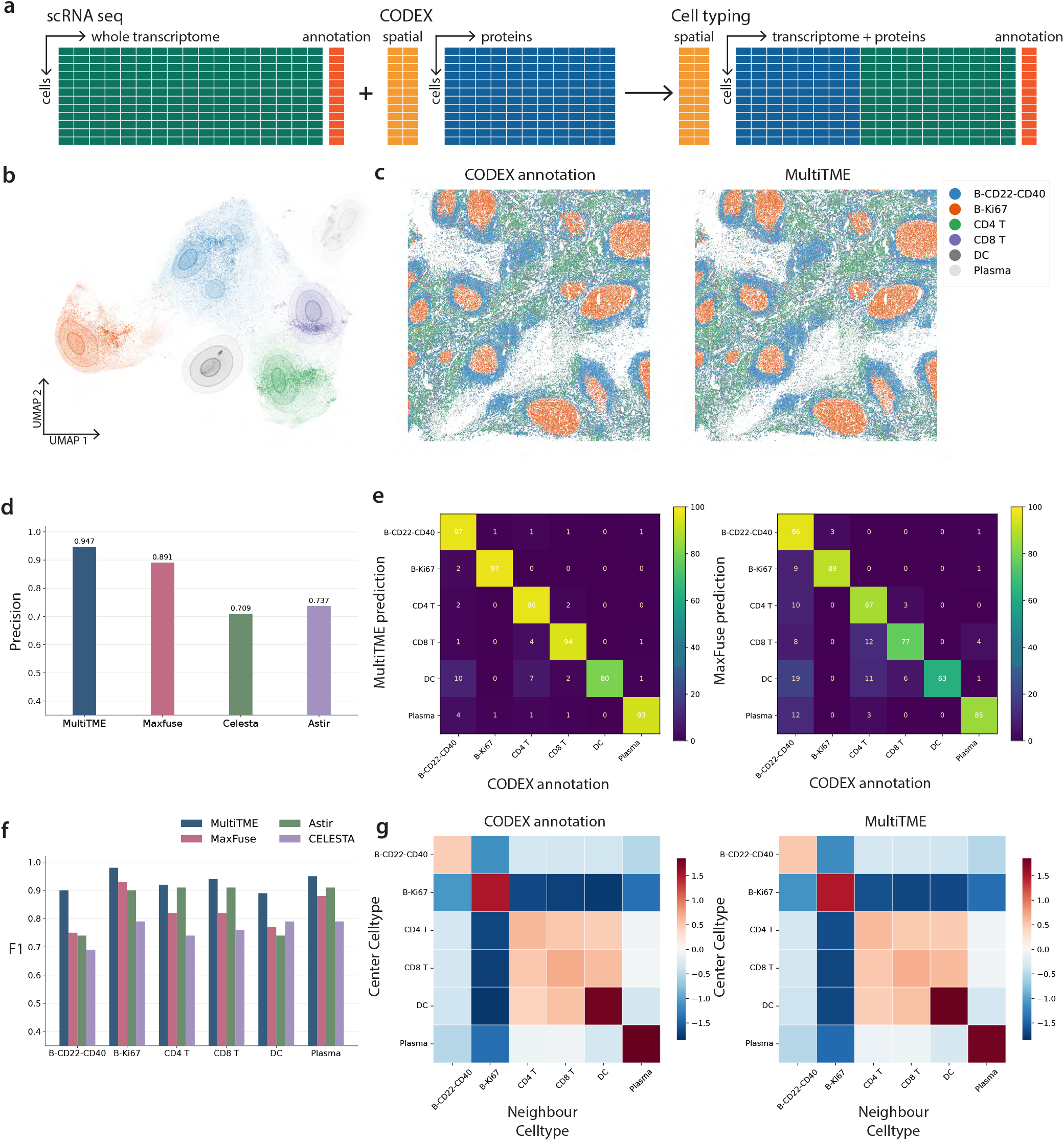
MultiTME enables spatially guided cell typing in spatial multimodal data. **(a)** Schematic of the cell typing task. **(b)** UMAP visualization shows alignment of cellular states across modalities. **(c)** Spatial cell type maps of the tonsil tissue. Ground-truth CODEX annotations (left) and MultiTME predictions (right) show highly consistent spatial organization of major immune populations. **(d)** Overall cell typing accuracy comparison across methods. MultiTME achieves the highest accuracy (0.947), outperforming the cross-modal integration method MaxFuse (0.891) and proteomics-based classifiers Celesta (0.709) and Astir (0.737). **(e)** Confusion matrices comparing predicted and ground-truth cell types for MultiTME and MaxFuse. **(f)** cell type–specific F1 scores across methods. MultiTME achieves consistently higher accuracy across all six immune populations. **(g)** cell type neighborhood enrichment matrices derived from spatial data.

To benchmark cross-modal cell typing performance, we compared MultiTME against representative integration and cell-type classification methods, including Celesta ^20^ and Astir ^21^ for proteomic cell typing and MaxFuse ^22^ for cross-modal alignment. MultiTME achieves the highest overall cell typing accuracy (0.947), outperforming MaxFuse (0.891), Celesta (0.709), and Astir (0.737) (Figure 2d). MultiTME’s performance is robust across cell types, outperforming competing methods in all six annotated cell states (Figure 2f). Confusion matrix analysis shows that MultiTME reduces cross-lineage misclassification relative to MaxFuse, particularly among closely related lymphocyte populations such as CD4 and CD8 T cells and between germinal-center and proliferating B-cell states (Figure 2e).

We also investigated whether MultiTME performance improves the consistency of cell type assignments across spatial neighborhoods. By computing cell type neighborhood enrichment matrices, we noted the interaction patterns inferred from MultiTME predictions closely match those derived from the ground-truth CODEX annotations (Figure 2g), indicating that MultiTME preserves the spatial organization of immune cell populations. Predicted and true local cell type fractions also show strong agreement across spatial neighborhoods (Figure S4). These results demonstrate that MultiTME provides a unified framework for accurate, integrated cell typing in spatial proteomics with matched scRNA-seq.

### MultiTME accurately completes spatial transcriptomic panels from joint scRNA and Xenium embeddings

We next sought to test whether MultiTME could effectively merge single-cell and spatial transcriptomics data to improve spatial analyses. Imaging-based spatial transcriptomics platforms (e.g., MERFISH, ^23^ CosMx, and Xenium) require designing probe panels that typically cover only a subset of the transcriptome. Single-cell RNA-seq offers a complementary data modality, providing whole-transcriptome coverage but lacking spatial resolution. We hypothesized that joint modeling of scRNA-seq and imaging-based spatial transcriptomics with MultiTME would learn a shared latent space that would allow applying the whole transcriptome scRNA-seq decoder to the imaging-based spatial transcriptomics cells to impute missing genes in the spatial panel (Figure 3a).

**Figure 3:**
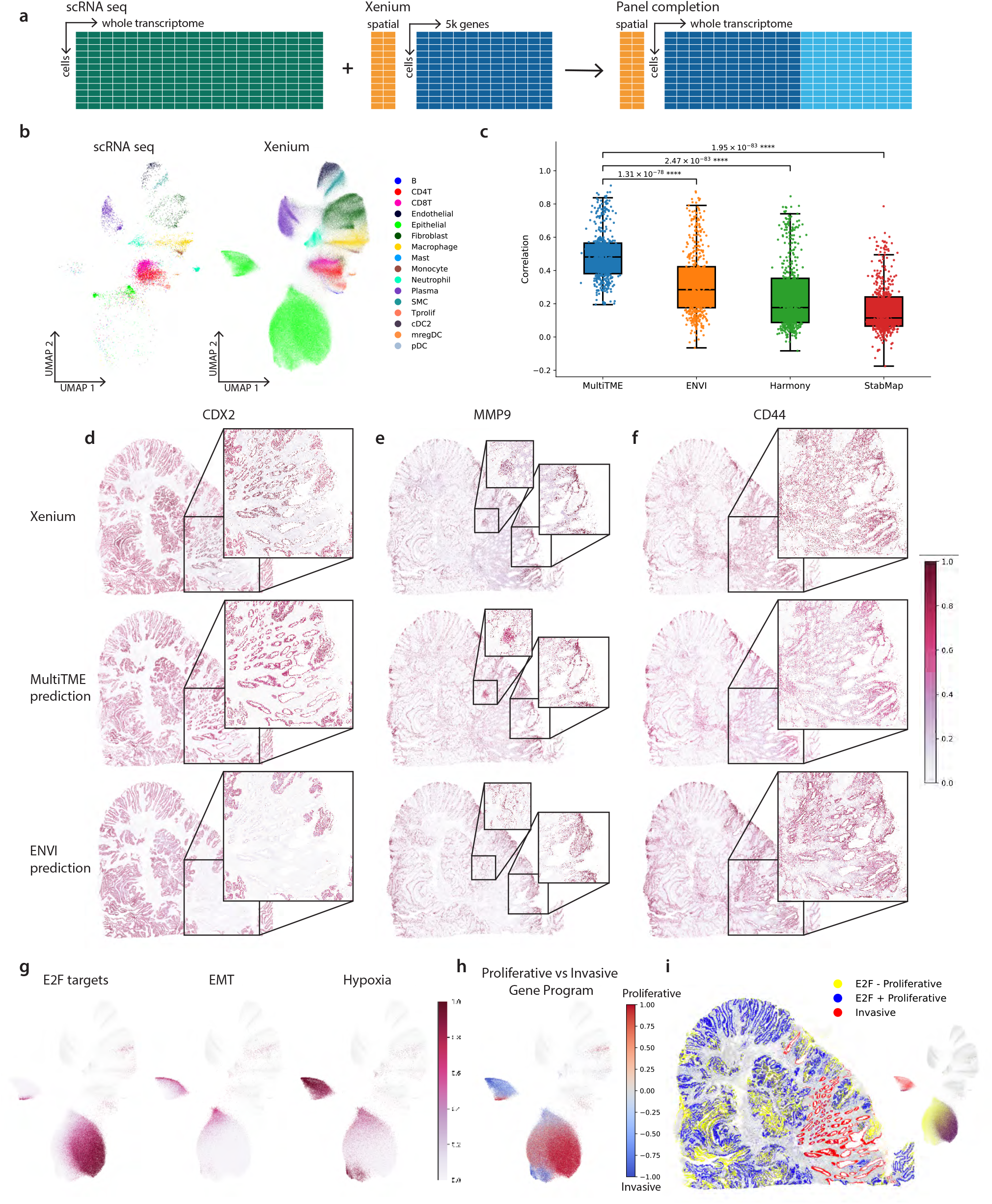
MultiTME accurately completes spatial transcriptomic panels from joint scRNA– Xenium embeddings. **(a)** Schematic of the panel completion task. **(b)** UMAP embeddings of the shared latent space learned by MultiTME, colored by cell type, shown separately for scRNA-seq (left) and Xenium (right). Concordant clustering across modalities indicates successful manifold alignment. **(c)** Per-gene Pearson correlation between predicted and ground-truth Xenium expression for MultiTME, ENVI, Harmony, and StabMap. *P* -values are from two-sided Wilcoxon signed-rank tests. **(d-f)** Spatial expression maps of *CDX2, MMP9*, and *CD44* in a colorectal cancer tissue section. Insets highlight regions where MultiTME more faithfully recovers the spatial expression pattern. **(g)** Tumor relevant Hallmark pathway enrichment scores using the completed transcriptomic panel. **(h)** Marker genes for invasive vs proliferative tumor projected onto the Xenium UMAP manifold. **(i)** Spatial distribution of *E2F* + proliferative, *E2F* - proliferative, and invasive tumor cells.

We trained MultiTME on a colon adenocarcinoma dataset ^6^ with scRNA-seq and Xenium data from the same patient. The joint embedding learned by MultiTME successfully aligned scRNA-seq and Xenium manifolds while preserving biologically meaningful structure (Figure 3b). Major lineages, including epithelial, T cells (CD4T, CD8T, Tprolif), B cells, myeloid (macrophage, monocyte, mregDC, cDC2, pDC), and stromal populations (fibroblast, endothelial, smooth muscle), overlap across modalities rather than forming assay-specific clusters, suggesting that the cyclic objective captures cell state variation shared between dissociated and in situ measurements.

To evaluate MultiTME’s ability to complete gene panels, we encoded the observed Xenium cells into the latent space using the trained Xenium panel encoder and decoded it back using the scRNA-seq decoder, thereby imputing the full transcriptome at single-cell resolution. To benchmark imputation accuracy, we randomly held out 500 genes from the Xenium panel. After imputing the full transcriptome, we computed the per-gene Pearson correlation between each heldout gene’s predicted and observed expression across cells. We compared MultiTME performance with ENVI ^12^, Harmony ^24^, and StabMap ^25^. MultiTME achieved a median per-gene Pearson correlation of approximately 0.5, whereas ENVI, Harmony, and StabMap achieved significantly lower median correlations (Figure 3c; all pairwise comparisons *p <* 10^−78^, Wilcoxon signed-rank test). Notably, the improvement was not driven by a small number of well-predicted genes. The entire distribution of gene-wise correlations shifts upward relative to all baselines, indicating a comprehensive improvement in panel completion across the transcriptome (Figure S1).

Spatial maps of selected genes further highlighted the advantages of MultiTME over the best-performing baseline method, ENVI (Figure 3d-f). For the epithelial lineage marker *CDX2* ^26^, the MultiTME prediction accurately recapitulated the glandular epithelial regions observed in the ground-truth Xenium, while ENVI produced diffuse signals that failed to resolve several tumor glands (Figure 3d insets). For the invasion-and matrix-remodeling–associated gene *MMP9*, ^27^ MultiTME captured localized expression at the invasive front and within stromal pockets, whereas ENVI underestimated these hotspots and yielded a spatially blurred pattern (Figure 3e insets). Analogous trends were observed for *CD44*, a marker of cancer stem cells and immune activation ^28^, where MultiTME accurately predicted a weaker and more diffuse signal while ENVI produced overly localized regions with bias towards epithelial niches (Figure 3f insets). Across all three exemplars, the improved accuracy of the MultiTME prediction led to better preservation of fine-grained spatial heterogeneity and predictions that were more biologically aligned with the ground-truth signal.

### Joint embedding reveals a spatial proliferative–invasive tumor axis

Beyond accurate gene-level completion, the integrated latent space learned by MultiTME organized tumor cells along a structured proliferative–invasive axis (Figure 3g, h). Projection of Hallmark pathway scores onto the Xenium embedding revealed that the two epithelial clusters correspond to distinct cell states within the cell type. Regions with high *E2F* activity exhibited low *EMT* and hypoxia signaling, while areas enriched for *EMT* and hypoxia displayed reduced proliferative signatures. Additional canonical colorectal cancer pathways also emerged with biologically expected cell type specificity (Figure S2). WNT/*β*-catenin signaling was enriched in epithelial/tumor states with minimal signal in immune lineages (Figure S2a), consistent with the canonical role of WNT pathway activation in colorectal carcinogenesis ^29^. MYC Targets V1 and V2 activity concentrated in the epithelial/tumor compartment and in proliferating T cells (Figure S2b-c), consistent with a tumor-intrinsic and proliferation-associated MYC program. On the other hand, invasive pathways such as TGF-*β* and glycolysis ^30,31^ are upregulated in the stress-adapted, invasive program.

Based on this mutually exclusive expression pattern, we designed a composite proliferative–invasive gene program that contrasts proliferation-associated genes (e.g., *LGR5, MKI67, TOP2A, CCNB1, RRM2*) against invasion-associated genes (e.g., *ZEB2, ITGA5, SERPINE1, VEGFA, SLC2A1, LDHA, PFKFB3, ICAM1*). This program reflects the canonical “go versus grow” tradeoff ^32^, in which tumor cells partition between proliferative expansion and invasive remodeling states ^33^. In the learned latent space, this program yielded a continuous gradient that separated proliferative from invasive tumor cells (Figure 3h). Spatial mapping of tumor cells using the composite proliferative–invasive score demonstrated spatially-distinct transcriptional states (Figure 3i). Proliferative regions corresponded to glandular epithelial compartments characterized by elevated *E2F* signaling, whereas invasive regions aligned with stromal interfaces, featuring activation of EMT, hypoxia, and matrix remodeling programs.

The shared embedding captured not only cell type identity but also tumor-state transitions. The proliferative–invasive axis generated important cell-state variation within epithelial populations, suggesting that cross-modal integration enables the recovery of functional tumor plasticity in a spatial context. Moreover, this proliferative–invasive differentiation is not identifiable from the original Xenium data (Figure S3). In colorectal cancer, tumors enriched for mesenchymal and invasive transcriptional states correspond to the CMS4 consensus molecular subtype ^34^, which is associated with worse relapse-free and overall survival. The ability to spatially resolve this axis at cellular resolution within intact tissue could inform strategies for targeting the invasive front, where therapy-resistant, EMT-high tumor cells interface with the stromal microenvironment. Direct UMAP embeddings derived from Xenium fail to align with the scRNA and separate among cell types (Figure S3a), and pathway activity computed from the restricted gene set does not reveal a clear distinction between proliferative and invasive tumor cells (Figure S3d-f). The unique biological insight enabled by MultiTME highlighted the power of merging complementary transcriptomic technologies and imputing whole transcriptomes onto spatial panel data.

### MultiTME maps spot transcripts onto CODEX cells for joint mRNA-protein spatial profiling at single cell resolution

A key limitation of high-resolution spatial proteomic platforms such as CODEX^35^ is their restricted panel size, typically measuring tens of proteins per cell. In dissociated cells, CITE-seq pairs proteomic measurements with whole transcriptome scRNA-seq to enable comprehensive multimodal single-cell readouts that enable better interpretation of cell state. In spatial transcriptomics, spot-based assays capture genome-wide transcriptomes but at multi-cellular spot resolution, obscuring cell-level heterogeneity. We hypothesized that MultiTME could bridge these complementary transcriptomic and proteomic spatial modalities, enabling the assignment of whole-transcriptome expression profiles to individual CODEX-resolved cells.

We applied MultiTME to the same colorectal cancer tissue as the previous section, but now focusing on different slices profiled by Visium HD and CODEX on consecutive tissue slices. ^6^ To evaluate whether the assigned transcripts faithfully captured spatial gene expression patterns, we constructed a synthetic Visium standard-definition (SD) dataset by coarsening the Visium HD bins. The original Visium HD was treated as a high-resolution ground truth and not used in training. MultiTME then jointly embedded spot-level transcriptomic measurements from the Visium SD data and cell-level proteomic measurements into a shared representation. Full transcriptomic profiles were then decoded onto each CODEX cell (Figure 4a).

**Figure 4:**
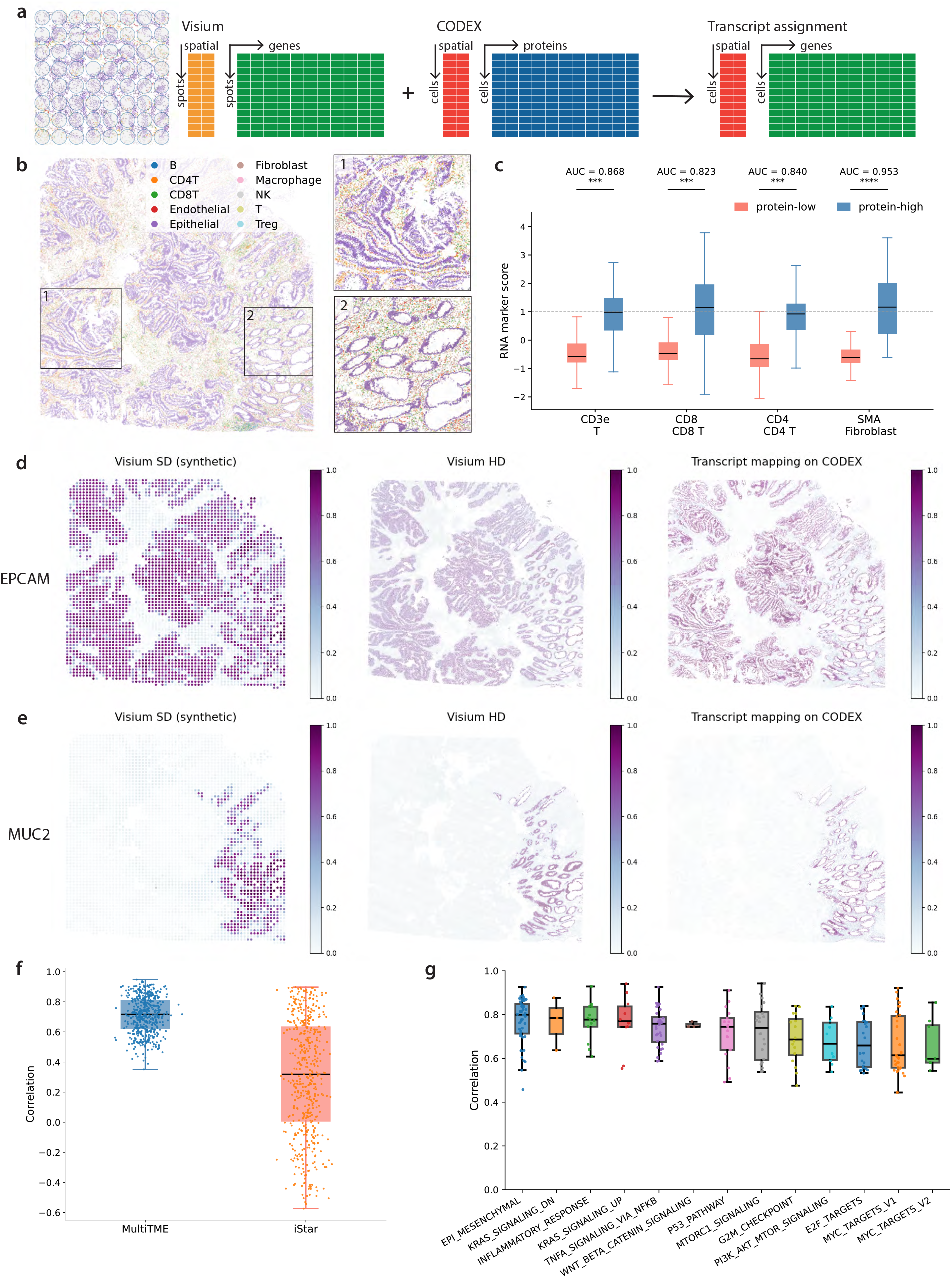
MultiTME assigns whole-transcriptome profiles to single cells by jointly modeling Visium and CODEX. **(a)** Schematic of the transcript assignment task. **(b)** Cell type map of a colorectal cancer tissue section with cell type annotation. **(c)** Concordance between measured protein abundance and MultiTME-assigned transcripts, including CD3e-marked T cells, CD8-marked cytotoxic T cells, CD4-marked helper T cells, and SMA-marked fibroblasts. AUC values indicate discrimination of protein-high versus protein-low cells by the matched RNA marker score. **(d-e)** Spatial expression maps of *EPCAM* (**d**) and *MUC2* (**e**) shown across three representations: synthetic Visium SD (left), Visium HD ground truth (center), and MultiTME transcript mapping on CODEX (right). **(f)** Per-gene Pearson correlation between predicted transcript assignments and Visium HD ground truth for MultiTME and iStar. **(g)** Per-pathway Pearson correlation between Hallmark gene set enrichment scores computed from MultiTME transcript assignments and from Visium HD, across 14 cancer-relevant pathways.

The resulting cell type map recovered the expected spatial organization of the colorectal tumor microenvironment (Figure 4b). Epithelial and tumor cells were mapped as lining the glandular structures, while immune cells, including CD4T, CD8T, B cells, NK cells, Treg, and macrophages, together with fibroblasts and endothelial cells, were observed in the surrounding stroma. For *EPCAM*, an epithelial adhesion molecule, the MultiTME transcript mapping onto single cells was necessary to resolve the ductal tissue architecture visible in Visium HD (Figure 4d). For *MUC2*, a goblet cell mucin, MultiTME similarly recovered the expected restriction to secretory epithelial cells within glandular lumina, preserving the spatial pattern observed in Visium HD at a resolution unattainable from the original Visium SD spots (Figure 4e). Notably, the MultiTME result appeared sharper than even the Visium HD, likely due to Visium HD’s bin-based quantification. By projecting transcripts onto individually segmented CODEX cells, MultiTME recovered crisper boundaries between epithelial glands and the surrounding stroma.

We next sought to quantitatively evaluate the MultiTME transcript mappings. To validate the biological fidelity of the assigned transcripts, we compared MultiTME-derived RNA marker scores against independently measured protein markers in the same cells. Cells with high protein abundance for CD3e, CD8, CD4, or SMA showed enriched matched RNA marker programs relative to protein-low cells, corresponding to T cells, CD8+ cytotoxic T cells, CD4+ helper T cells, and activated fibroblasts, respectively (Figure 4c). The RNA marker scores accurately distinguished protein-high from protein-low cells across all programs, with AUC values derived from Mann-Whitney U test statistics ranging from 0.823 to 0.953, supporting preservation of expected protein-RNA concordance. These AUC values represent the probability that a randomly selected protein-high cell has a higher matched RNA marker score than a randomly selected protein-low cell, providing a sample-size-invariant measure of concordance between independently measured protein and predicted transcript programs. We also compared the transcript assignment with iStar ^36^, a multimodal method for histology-assisted super-resolution in spatial transcriptomics. We aggregated MultiTME and iStar predictions, along with Visium HD ground truth, into 50*µm* ×50*µm* bins to enable comparison at a common spatial resolution, then computed Pearson correlation between predicted and measured expression per gene across bins (Methods). MultiTME outperformed iStar in per-gene correlation between predicted and ground-truth expression (Figure 4f), achieving a median gene-wise Pearson correlation of approximately 0.7, compared with approximately 0.1 for iStar.

Beyond individual genes, we assessed whether the assigned transcriptomes preserve pathway-level biological programs by computing Hallmark gene set enrichment scores for each cell and correlating these with scores derived from Visium HD (Figure 4g). Across a panel of 14 cancer-relevant pathways, MultiTME consistently achieved high per-pathway correlations (median ∼0.75–0.85). Tumor-intrinsic programs such as MYC Targets V1/V2, E2F Targets, G2M Checkpoint, MTORC1 Signaling, and PI3K/AKT/MTOR Signaling were consistently recovered, as were microenvironmental signatures including Inflammatory Response, TNF*α* Signaling via NF-*κ*B, and Epithelial-Mesenchymal Transition. Signaling pathways that are important to colorectal carcinogenesis, like KRAS Signaling Up and Down, WNT/*β*-catenin Signaling, and the P53 Pathway, were also accurately assigned at single-cell resolution. The uniformly high correlations across functionally diverse gene sets suggest that MultiTME’s transcript assignment preserves higher-order transcriptional programs rather than merely fitting individual marker genes.

These results demonstrate that MultiTME can leverage the complementary strengths of spot-level transcriptomics and cell-level proteomics to produce spatially resolved, multimodal spatial profiles at single-cell granularity. Protocols exist for performing post-Visium HD spatial proteomics on the same slice. In principle, this could obviate the need for unpaired merging and SD integration. In practice, the high cost and relatively smaller field of view of Visium HD and other fine-grained spot-based spatial transcriptomics platforms presents limitations to scaling such a strategy. Technical limitations with tissue warp and degradation between assays also make consecutive tissue slices a pragmatic alternative in many scenarios. Finally, MultiTME goes beyond simply increasing spot resolution: it provides direct mapping of transcripts onto individual cells to merge unpaired modalities into paired, multimodal spatial profiles at single-cell resolution.

### Modality translation reveals and corrects platform-specific technical biases

Imaging-based spatial transcriptomic platforms all function on the same fundamental concepts (i.e. engineered probes, in situ hybridization) but differ in technical details of their implementations. These implementation differences lead to technology-specific biases that confound biological results. With new studies emerging rapidly, meta-analyses and large pan-tissue/disease studies are soon to follow that will necessitate integrating imaging-based spatial transcriptomics data across multiple platforms. We hypothesized that MultiTME could disentangle or remove these platform-specific batch effects and thereby harmonize across technologies to enable effective integration across platforms. To test this, we evaluated MultiTME on five tissues from a recent benchmark dataset ^6^, comparing CosMx 6K and Xenium 5K on serial slices.

We first quantified baseline gene expression correlations across the two platforms across the five tissues. For each tissue, we computed pseudobulk profiles by averaging expression within each of 10 cell types per platform for comparison (Methods). Direct comparison of raw measurements revealed only modest concordance with a linear regression slope *b* = 0.423 (Figure 5a, left), with substantial dispersion away from the identity line. Notably, several genes appeared over-detected in CosMx relative to Xenium, consistent with background-driven inflation of low-level signals. ^37^

**Figure 5:**
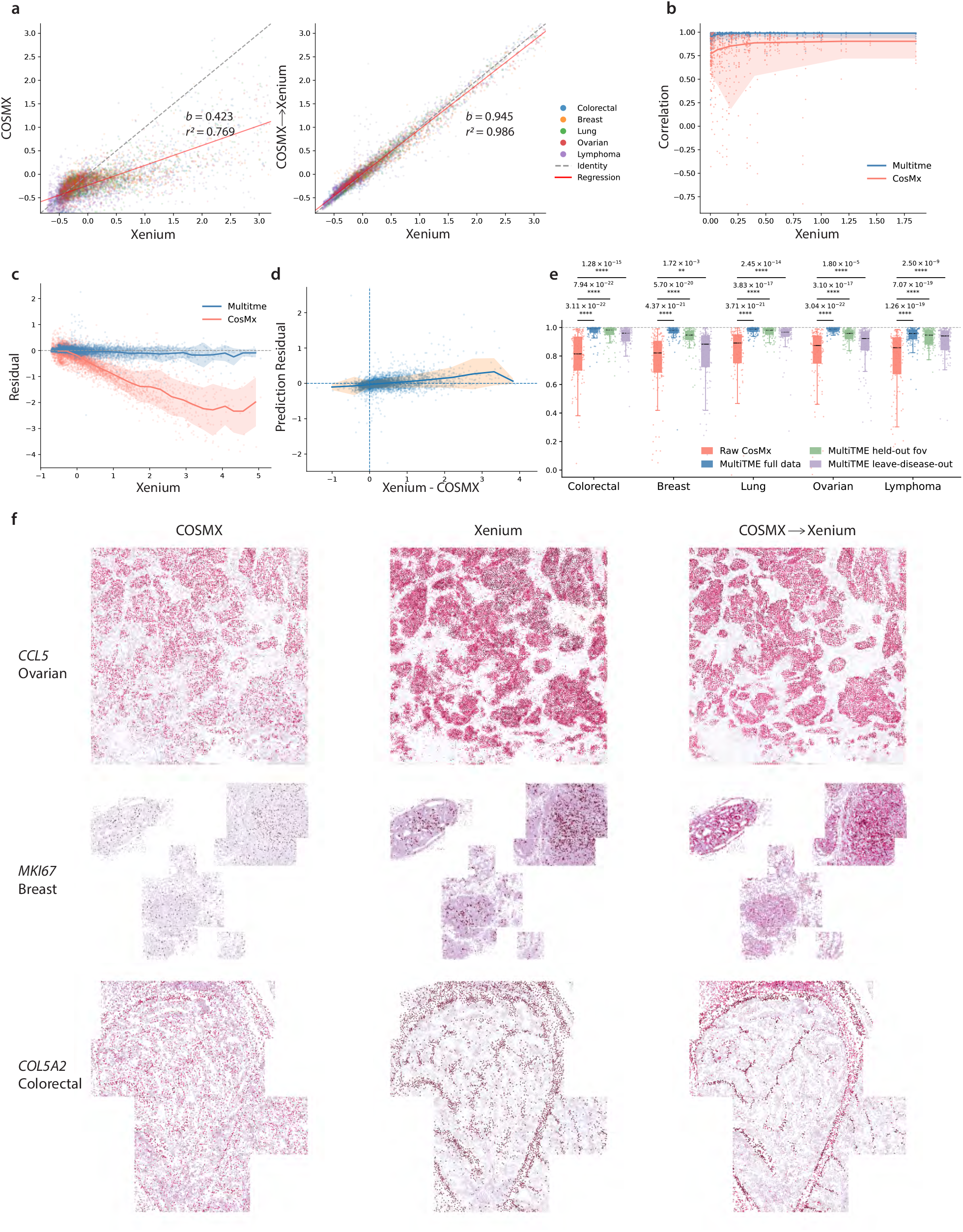
MultiTME corrects modality-specific biases across spatial transcriptomics platforms. **(a)**Raw cross-platform measurements show substantial dispersion from the identity line, whereas MultiTME translation from CosMx to Xenium improves concordance. **(b)** Gene-wise correlation across expression levels is consistently higher after translation. (**c–d**) Prediction residual is centered near zero and remains stable both across raw Xenium expression and across raw Xenium–CosMx discrepancy. (**e**) Per-gene correlation distributions across five cancer types demonstrate improved cross-platform agreement. (**f**) Representative spatial examples show that translation suppresses diffuse background and restores sharper spatial expression patterns across genes and tissue types.

MultiTME was then fit to each pair of serial sections from the same tissue. After translation from CosMx to Xenium through the MultiTME latent space, the agreement significantly improved, with the fitted relationship approaching the identity line and the coefficient of determination increasing substantially (Figure 5a, right). This improvement was consistently higher across expression levels, with the greatest gains observed for low-and moderately expressed genes (Figure 5b). The prediction residual remains consistently centered near zero, indicating that MultiTME does not introduce systematic bias as expression level changes (Figure 5c). Likewise, the residual remains stable against raw Xenium–CosMx difference, showing that the translated predictions are robust even when the two platforms initially disagree substantially (Figure 5d). This suggests that MultiTME is particularly effective at correcting background-driven distortions that disproportionately affect weak signals.

Spatial examples illustrated the biological effect of this correction (Figure 5f). For *CCL5* in ovarian cancer, *MKI67* in breast cancer, and *COL5A2* in colorectal cancer, raw CosMx measurements appeared more diffuse and background-contaminated, whereas Xenium exhibited sharper and more localized spatial structure. Translation from CosMx to Xenium suppressed this diffuse background while preserving the major anatomical and microenvironmental patterns.

To assess robustness across disease contexts, we summarized per-gene cross-platform correlations for five cancer types: colorectal, breast, lung, ovarian, and lymphoma ^37^. In every dataset, MultiTME translation improves agreement relative to raw Xenium measurements (Figure 5e). To test generalization, we further evaluated two generalization settings: predictions on field-of-view (FOV) regions excluded from training within the same cancer type, and a leave-one-disease-out setting in which the target cancer type was entirely withheld during training. Held-out FOV performance was comparable to the full data setting across all five tissues, confirming that MultiTME is robust within a tissue type and generalizes across spatial regions within a tissue. The leave-disease-out setting was more challenging, requiring MultiTME to extract generalizable technical translation strategies applicable across distinct tissues. As expected, performance in this setting was worse than in the matched tissue type settings. However, even in this case MultiTME consistently outperformed the original CosMx signal across all tissues. These results demonstrate that MultiTME can filter out assay-specific artifacts to capture latent biological states that generalize across tissues.

## Discussion

Cross-modal integration in spatial biology is fundamentally constrained by the lack of paired measurements. While existing approaches perform well when modalities share strongly linked features, many emerging datasets—such as combinations of targeted spatial assays and whole-transcriptome sequencing—exhibit weak or indirect correspondence. In these settings, integration requires recovering a shared biological structure without relying on explicit feature matching.

MultiTME is motivated by this setting and addresses it by learning a cycle-consistent shared latent representation. By enforcing that cross-modal translations preserve both latent states and observable features, the model constrains mappings between modalities to be mutually informative. This enables stable transfer of information across assays, even when feature spaces differ substantially. As a result, the learned representation supports not only alignment but also reconstruction of unmeasured modalities. This cycle-consistent approach enabled MultiTME to achieve state-of-the-art accuracy in cross-modal cell typing, transcriptome panel completion, and multimodal superresolution.

MultiTME mappings are dependent on the biological signal in the input datasets and the quality of any optional expert annotations. The semi-supervised cell typing component assumes scRNA-seq annotations are correct when transferring labels to other modalities. Likewise, any curated proteomic markers used to anchor spatial cells are assumed to be accurate identifiers of their corresponding cell types for the highest expressing cells. Upstream spatial preprocessing issues, such as segmentation errors and serial slice misalignment, can also dilute or confound biological signal. Extensions to MultiTME that model cells in spatial transcriptomics as admixtures rather than individual cells, analogous to recent unimodal methods like SPLIT, ^38^ may enable MultiTME to overcome these preprocessing issues to further boost performance.

The ability of MultiTME to unify across technical platforms, as we demonstrated in Figure 5, presents the potential to serve as the basis for an approach to multimodal atlas integration and harmonization. The ability of MultiTME to scale to tens or hundreds of datasets is still unclear, and the theoretical and empirical limits of the cycle-consistency strategy in MultiTME have not yet been established. It is possible that fine-tuning of the model architecture or loss functions may be necessary to adapt to integrating hundreds of disparate datasets in pan-tissue and pan-disease studies. Nevertheless, with new single-cell and spatial datasets rapidly emerging, the core framework underlying MultiTME holds great promise as a general-purpose tool for cross-dataset multimodal studies. We anticipate that MultiTME and future versions will be particularly useful in multimodal atlas construction, where the key challenges are integrating and harmonizing across disparate, unpaired modalities while preserving biological structure, thereby enabling novel scientific discoveries.

## Methods

MultiTME is a variational autoencoder that maps observations from multiple spatial omics platforms into a shared latent space. The model takes as input a set of modality configurations specifying the feature dimensionality and likelihood family for each platform. Let *M* denote the set of modalities, *D*_*m*_ the feature dimension of modality *m*, and 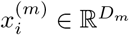 the observed feature vector for cell *i* in modality *m*. The model assumes that all observations arise from a shared latent variable *z*_*i*_ ∈ℝ^*d*^ (*d* = 32) that captures the underlying biological state of the cell.

### Model architecture

MultiTME follows a variational autoencoder architecture with modality-specific projection layers, a shared encoder that maps all modalities to a common latent space, and modality-specific decoders for reconstruction.

For *M* modalities with feature dimensions *D*_1_, *D*_2_, …, *D*_*M*_, each modality *m* is first mapped to a common intermediate dimension *h* through a modality-specific projection layer

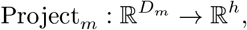

implemented as a linear layer followed by layer normalization and ReLU activation. In practice, we set *h* = 256.

The projected representation is then processed by a shared encoder network

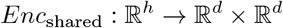

that outputs the mean and log-variance of a Gaussian variational posterior

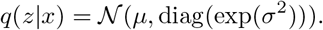

The shared encoder consists of a single hidden layer (128 units with layer normalization and ReLU) followed by separate linear heads producing *µ* and log *σ*^2^. The log-variance is clamped to the range [−4, 4] for numerical stability.

The overall modality-specific encoder can therefore be written as

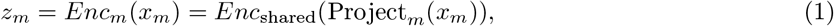

Where 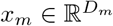 is the observation from modality *m*.

Importantly, the shared encoder contains no modality identifiers or modality-specific parameters. Modality-specific transformations occur only in the projection layers, which allows the model to accommodate heterogeneous feature spaces (e.g., tens of proteins in CODEX versus thousands of genes in transcriptomic assays) while encouraging the latent representation to capture modality-invariant biological structure.

Each modality has a dedicated decoder network *Dec*_*m*_ that reconstructs observations 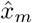 from the latent representation *z*_*m*_. Decoders follow the likelihood models appropriate for each modality (negative binomial for count-based assays and Gaussian for protein intensities).

#### Cycle consistency constraints

To align latent representations across different modalities without paired data, we enforce bidirectional cycle consistency. For each pair of modalities (*m*_1_, *m*_2_), we construct two cycle paths. For the forward cycle (*m*_1_ →*m*_2_), we encode 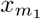 to obtain 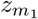 and decode it to predicted 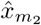 using 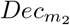. The 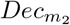 is then re-encoded to obtain 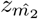 and decode to 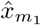 by 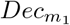. We also define the reverse cycle (*m*_2_ →*m*_1_) in the same fashion but opposite direction (Figure 1b). Cycle consistency is enforced at both the latent and observation levels through the following loss terms:

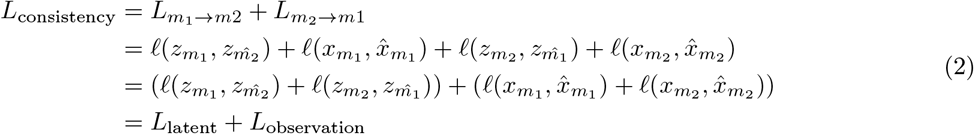

#### Spatial regularizer

An observation of spatial profiling data is that modalities acquired from consecutive tissue slices preserve similar spatial organization of cell types, even though individual cells are not directly matched across modalities.

For a spatial pair of modalities (*A, B*) measured on co-registered tissue sections, the alignment loss encourages cells that occupy the same spatial location and belong to the same cell type to have similar latent representations.

For each cell *i* in modality *A* and each candidate cell type *t*, we identify the *K* = 10 nearest neighbors of *i* in modality *B* using Euclidean distance in physical tissue coordinates. Let 𝒩_*B*_(*i*) denote this neighbor set. We compute a spatially weighted and type-weighted mean latent representation of these neighbors

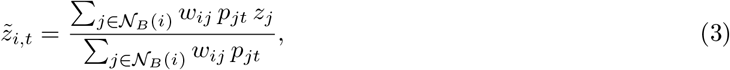

where *z*_*j*_ is the latent representation of neighbor cell *j, p*_*jt*_ is the probability that cell *j* belongs to type *t*, and *w*_*ij*_ is a spatial weight defined by a Gaussian kernel

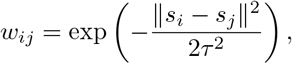

with *s*_*i*_ denoting spatial coordinates and *τ* = 100 *µm* controlling the spatial bandwidth.

The alignment loss minimizes the squared distance between the latent representation of cell *i* and the local cell-type target

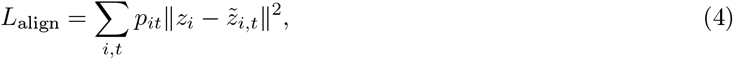

where *p*_*it*_ represents the probability that cell *i* belongs to type *t*. This weighting ensures that the loss activates only for cell types that are locally present.

#### Semi-supervised celltype regularizer

Cell-type probabilities *p*_*it*_ are determined through a three-tier hierarchy.

##### Annotated cells

For cells with known annotations (e.g. scRNA), smoothed one-hot labels (smoothing factor 0.05) are used as fixed targets with no gradient flowing through the classifier.

##### Marker-supported cells

For unlabeled cells in modalities with curated marker panels (e.g. CODEX), we employ a semi-supervised marker-based pseudo-labeling strategy. For each cell type *c*, we identify high-confidence exemplar cells by selecting those with the highest expression of known marker genes or proteins. For example, cells with high *CD4* expression are labeled as CD4^+^ T cells, while cells with high *Ki67* expression are labeled as proliferating cells. These pseudo-labeled cells provide weak supervision through a classifier attached to the latent representation.

##### Unlabeled cells without markers

For modalities without marker panels, cell-type probabilities are taken directly from the classifier softmax output.

#### Spatial tiling with halo exchange

Spatial mini-batches are constructed using a tiling strategy to preserve local spatial context during training. The shared spatial domain is partitioned into square tiles (default side length 500*µm*). Each training batch contains all cells whose coordinates fall within the tile interior (core cells), augmented with neighboring cells within a 150 *µm* border expansion (halo cells).

Halo cells are included only for spatial neighborhood computations but excluded from reconstruction and classification losses using Boolean masks. This prevents double-counting cells that appear in overlapping tiles while ensuring that spatial regularization always uses true physical neighbors.

This design follows the halo-exchange pattern used in domain-decomposition methods for numerical PDE solvers, where ghost cells provide boundary information for local computations without contributing independent updates.

## Metrics

### Comparison with iStar

To compare MultiTME and iStar at a common resolution, we constructed a regular 50*µm* ×50*µm* grid spanning the tissue and aggregated both MultiTME’s cell-based prediction and iStar’s pixel-based prediction into bins by summing the predicted expression of all source spots whose centroids fell within each bin (nearest-bin assignment via KD-tree). Visium HD ground truth was aggregated identically. For each gene, we computed the Pearson correlation between predicted and ground-truth expression across all bins. The bin size of 50*µm* ensures each bin aggregates sufficient Visium HD reads for stable correlation estimates.

### Pseudobulk comparison

For each platform and tissue, cells were assigned to one of 10 cell types provided in the celltype annotation. Pseudobulk profiles were computed by averaging expression across cells within each cluster for all genes included in the Xenium panel. To evaluate cross-platform concordance, we compared the distributions of per-gene correlations under raw and translated expressions using the Wilcoxon signed-rank test, with genes paired across conditions. We also fit an ordinary least-squares regression to the pooled (gene, cell-type) values and report *r*^2^ as a summary of overall fit.

## Code availability

All MultiTME source code is available from GitHub (https://github.com/tansey-lab/multitme).

## Data availability

All data analyzed in this study are publicly available from previously published sources. The scRNA-seq and CODEX tonsil data were obtained from Kennedy-Darling et al. ^19^; the colorectal cancer scRNA-seq, Xenium, Visium HD, and CODEX data were obtained from Ren et al. ^6^; and the multi-cancer CosMx–Xenium platform-comparison datasets were obtained from Cervilla et al. ^37^ Accession numbers and download links are provided in the corresponding original publications.

## Acknowledgements

W.T. is supported by the NIH/NCI (R37 CA271186, U54 CA274492, P30 CA008748), Break Through Cancer, the Fund for Innovation in Cancer Informatics, the Cancer AI Alliance, the Tow Center for Developmental Oncology, and the Maurice Campbell Initiative at Memorial Sloan Kettering Cancer Center.

## Author Information

### Author contribution

H.Z. led the development and implementation of the model and the analysis of datasets. J.F.Q. provided software engineering support to develop the MultiTME package. W.T. oversaw the project and guided the development of the models and interpretation of the results. The Break Through Cancer Data Science TeamLab members reviewed the manuscript and contributed to internal discussions during the development of the method.

### Full list of Data Science TeamLab

Cheng-Zhong Zhang, Ethan Cerami, Franziska Michor, Nathalie Agar, Rameen Beroukhim, Ashley Kiemen, Atul Deshpande, Dimitri Sidiropoulos, Genevieve Stein-O’Brien, Leslie Cope, Rachel Karchin, Roy Elias, Kadir Akdemir, Linghua Wang, Charlie Whittaker, Stuart Levine, Andrew McPherson, Benjamin Greenbaum, Niki Schultz, Sohrab Shah, Elana Fertig, Renad Al Ghazawi, Nil Aygun, Archana Balan, Gerard Baquer, Vincent Butty, Nick Ceglia, Jennifer Chen, Yu-Chen Cheng, Andrew Cherniack, Kate Cho, Kyung Serk (Kevin) Cho, Chase Christenson, Simona Cristea, Yibo Dai, Simona Dalin, Ino de Bruijn, Charlie Demurjian, Henry Dewhurst, Huiming Ding, Tom Dougherty, Emma Dyer, Yuval Elhanaty, Jiayi Fan, Sasha (Alexander) Favorov, Andre Forjaz, Samuel Freeman, Alessandro Grande, Christopher Graser, Benjamin Green, Eliyahu Havasov, Jason Hwee, Afrooz Jahedi, Jiahui Jiang, Taisha Joseph, Saurabh Joshi, Nancy Jung, Luciane Kagohara, Jennifer Karlow, Jiaying Lai, Kevin Meza Landeros, Joshua Lau, Nora Lawson, Jake Lee, Hannah Lees, Matt Leventhal, Alex Ling, Lester (Yunzhou) Liu, Yang Liu, Yunhe Liu, Dmitrijs Lvovs, Valentina Matos Romero, Marcus Mendes, Jose Meza Llamosas, Thomas Mitchell, Hideaki Mizuno, Jacob Myers, Matthew Myers, Nataly Naser Al Deen, Siri Palreddy, Sergiu Pasca, Jeffrey Quinn, Sabahat Rahman, Kimal Rajapakshe, Shriya Rangaswamy, Greg Raskind, Shahab Sarmashghi, Manuel Schürch, Alyza Skaist, Alexander Solovyov, Haruna Tomono, Michael Toomey, Chris Tosh, Mesut Unal, Yann Vanrobaeys, Miriam Vines, Jun Wang, Meredith Wetzel, Marc Williams, Juan Xie, Lingqun Ye, Long Yuan, Syed Zaidi, Matthew Zatzman, Haoran Zhang, Bo Zhao, Peter Zheng, Feiyang Huang, Irika Katiyar, Elana Sverdlik.

