## Supplementary figures for "Cycle-consistent deep generative modeling unifies cellular states across unpaired spatial and single-cell modalities"

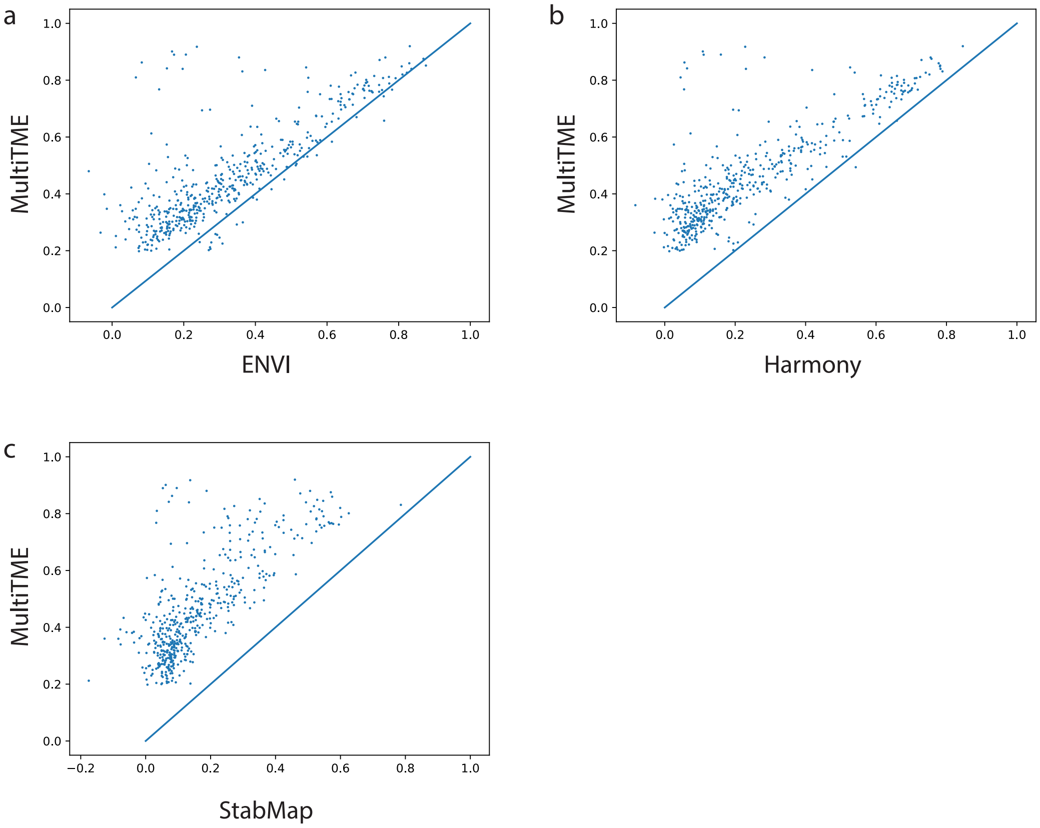


Figure S1: Scatter plots comparing per-gene Pearson correlations between MultiTME and each baseline method (ENVI, Harmony, StabMap) across 500 held-out genes. Each point represents one gene, and the diagonal line indicates equal performance. Points above the diagonal indicate genes where MultiTME achieves higher imputation accuracy. The distribution of gene-wise correlations shifts systematically above the diagonal across all baselines, indicating that MultiTME performs consistently better.

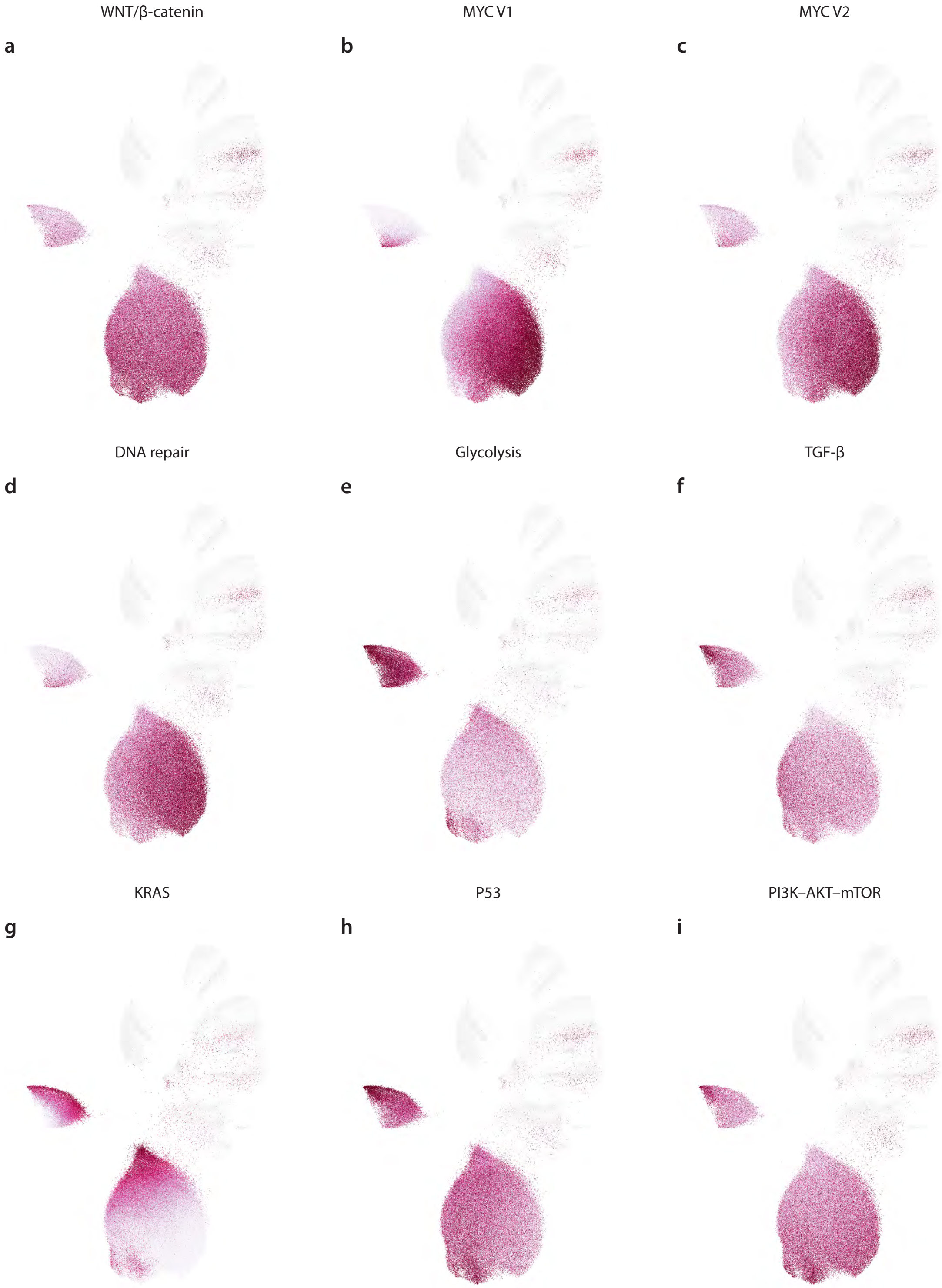
WNT/β-catenin
MYC V1
MYC V2
a
b
c
DNA repair
Glycolysis
TGF-β
d
e
f
KRAS
P53
PI3K–AKT–mTOR
g
h
i

Figure S2: (figue above) Spatial maps of additional pathway activity scores for selected cancer-associated signaling programs (WNT/$\beta$-catenin, MYC V1/V2, DNA repair, glycolysis, TGF-$\beta$, KRAS, p53, and PI3K–AKT–mTOR). Each panel shows the spatial distribution of cells colored by pathway activity (higher intensity indicates higher activity). Proliferation-associated pathways (MYC, DNA repair, WNT) share similar expression patterns as E2F that are upregulated in proliferative, stem-like tumor programs, whereas invasive pathways such as TGF-$\beta$ and glycolysis exhibit patterns like EMT that are upregulated in the invasive, stress-adapted program. Additionally, KRAS and PI3K–AKT–mTOR activity are more evenly distributed across tumor cells, suggesting that these pathways act as a shared oncogenic background.

Figure S3: (figure below) (a) UMAP visualization of Xenium and scRNA cells (subsets with the common genes that is observed in both Xenium and scRNA) shows significant modality separation. (b) Cell-type annotations further show that the scRNA occupies distinct regions from the Xenium cells, limiting direct cross-modality interpretation. (c–f) Pathway score calculated from the raw Xenium panel projections on the Xenium embedding. (c) The E2F program shows a broadly interpretable proliferative pattern, with enrichment in the major proliferative epithelial/tumor region. In contrast, (d) EMT, (e) hypoxia, and (f) TGF-$\beta$ programs are weakly resolved or sparsely detected, reflecting the limited gene coverage of the Xenium panel and insufficient pathway representation. These programs also fail to clearly separate the invasive cluster from the proliferative cluster, suggesting that reduced-panel Xenium measurements alone are insufficient to robustly distinguish these transcriptional states.

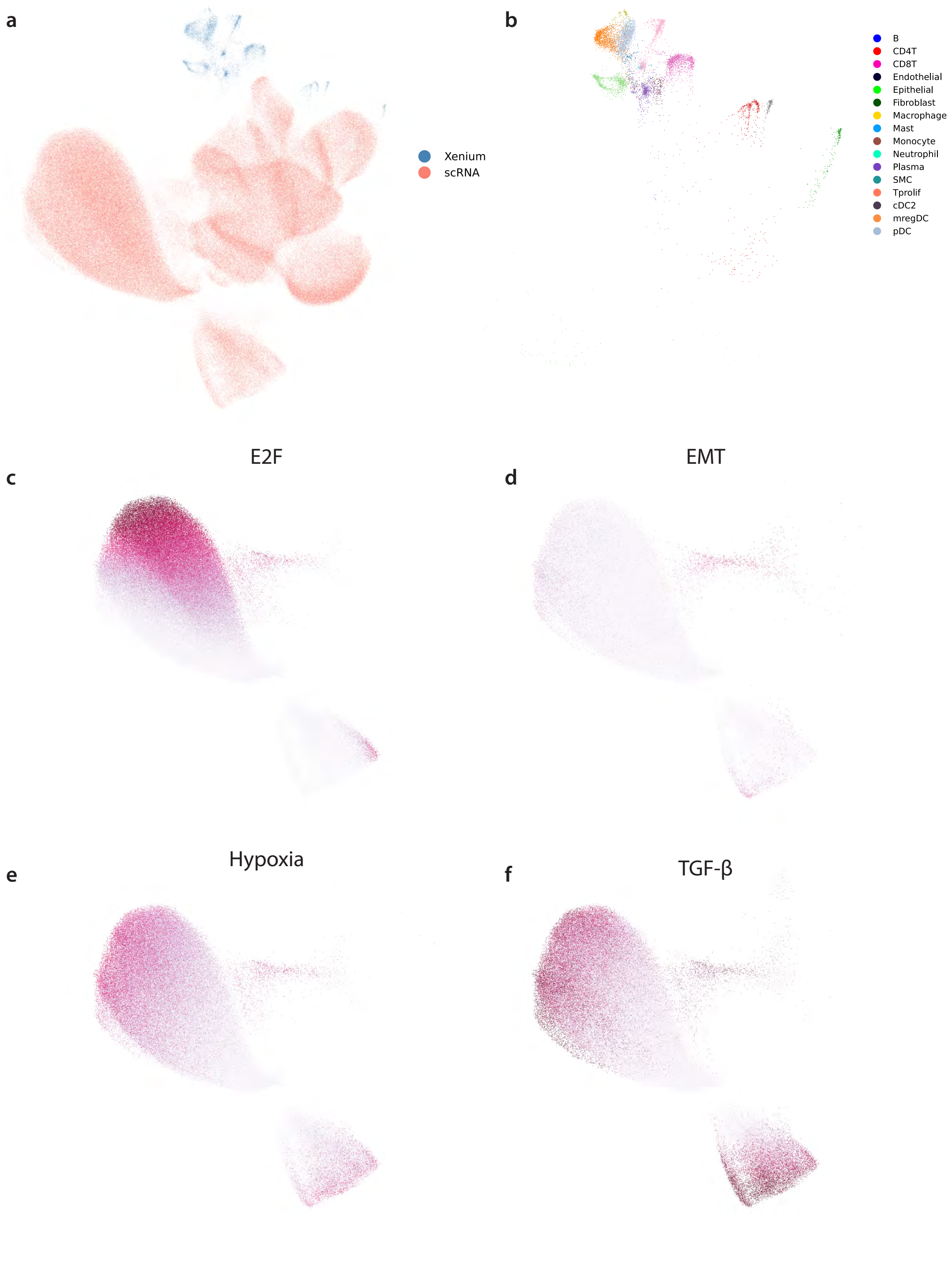
a
Xenium
scRNA
b
B
CD4T
CD8T
Endothelial
Epithelial
Fibroblast
Macrophage
Mast
Monocyte
Neutrophil
Plasma
SMC
Tprolif
cDC2
mregDC
pDC
c
E2F
d
EMT
e
Hypoxia
f
TGF-β

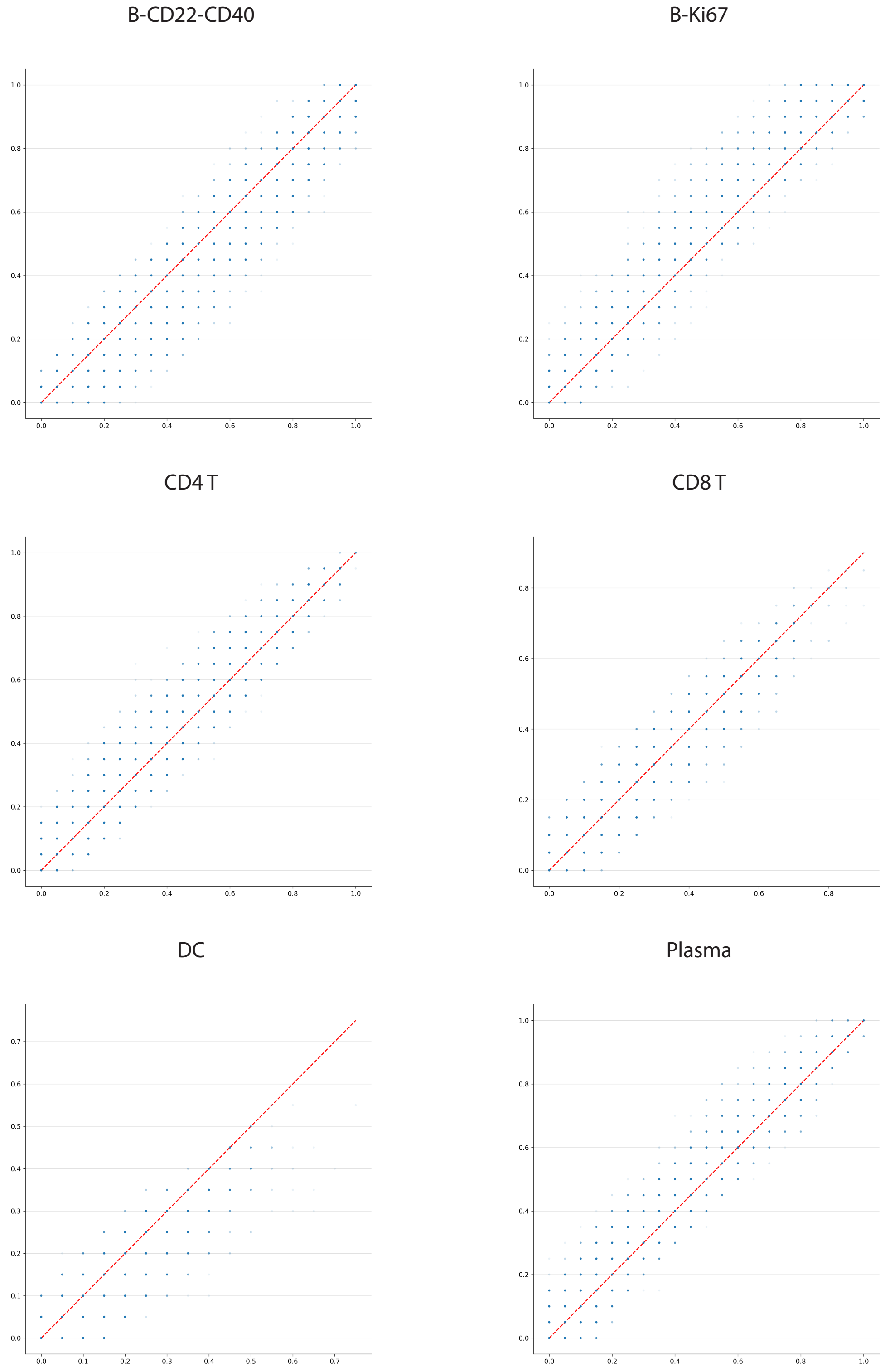


Figure S4: Scatter plots comparing predicted versus true local fractions of selected cell types across spatial neighborhoods (B–CD22–CD40, B–Ki67, CD4 T, CD8 T, dendritic cells (DC), and plasma cells). Each point represents a spatial neighborhood, with the x-axis denoting the true local fraction of a given cell type and the y-axis the corresponding model-predicted fraction. The dashed red line indicates the identity line ($y = x$). Predictions closely match ground truth for major lymphoid populations, while reduced agreement for dendritic cells likely reflects their lower abundance and increased variability in local composition.

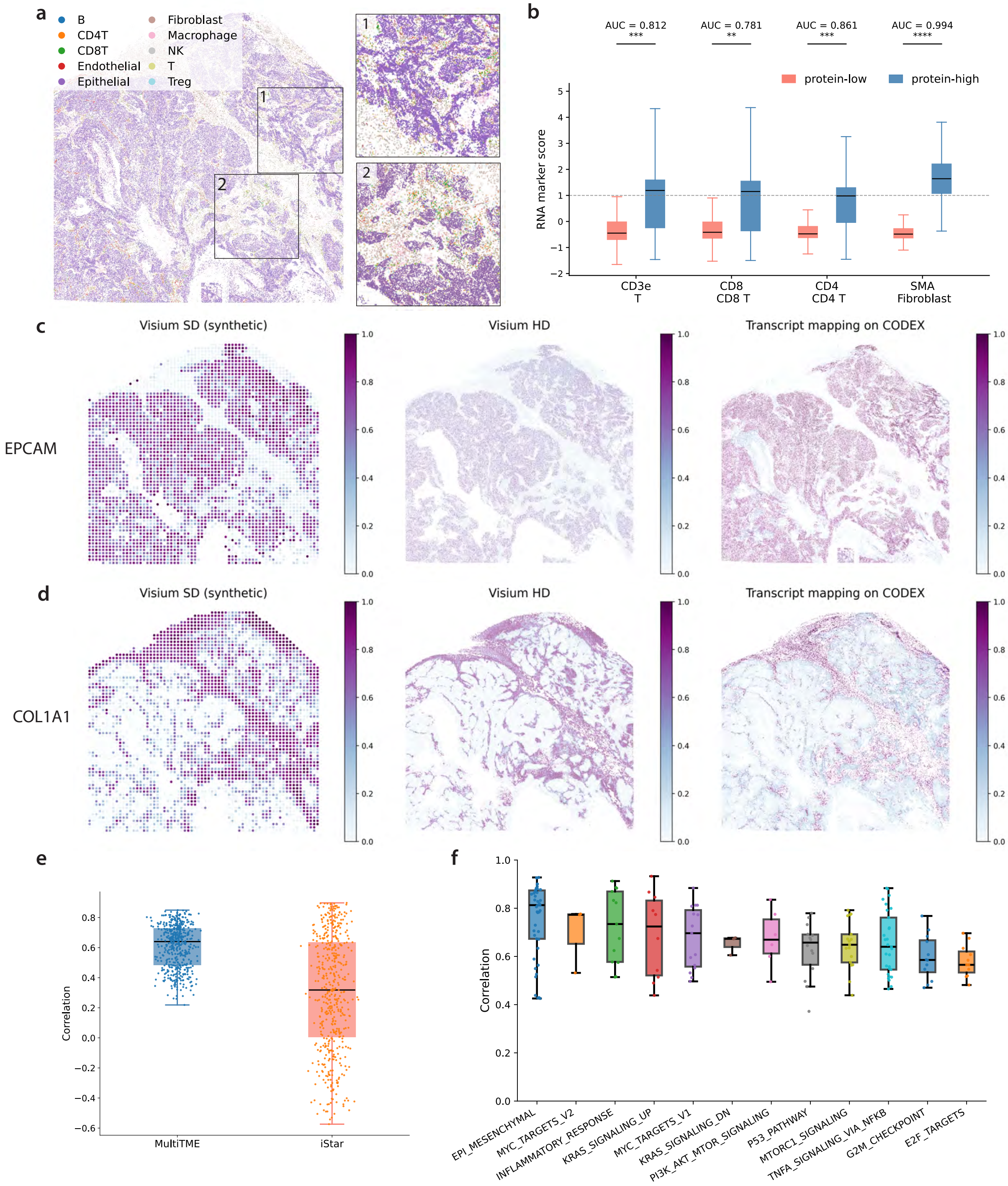
a
B
CD4T
CD8T
Endothelial
Epithelial
Fibroblast
Macrophage
NK
T
Treg
1
2
b
AUC = 0.812
***
AUC = 0.781
**
AUC = 0.861
***
AUC = 0.994
****
protein-low
protein-high
RNA marker score
CD3e
T
CD8
CD8 T
CD4
CD4 T
SMA
Fibroblast
c
Visium SD (synthetic)
Visium HD
Transcript mapping on CODEX
EPCAM
d
Visium SD (synthetic)
Visium HD
Transcript mapping on CODEX
COL1A1
e
Correlation
MultiTME
iStar
f
Correlation
EPI_MESENCHYMAL
MYC_TARGETS_V2
INFLAMMATORY_RESPONSE
KRAS_SIGNALING_UP
MYC_TARGETS_V1
KRAS_SIGNALING_DN
PI3K_AKT_MTOR_SIGNALING
P53_PATHWAY
MTORC1_SIGNALING
TNFA_SIGNALING_VIA_NFKB
G2M_CHECKPOINT
E2F_TARGETS

Figure S5: **MultiTME transcript assignment performance on ovarian cancer sample (a)** Cell type map of a ovarian cancer tissue section with cell type annotation. **(b)** Concordance between measured protein abundance and MultiTME-assigned transcripts, including CD3e-marked T cells, CD8-marked cytotoxic T cells, CD4-marked helper T cells, and SMA-marked fibroblasts. AUC values indicate discrimination of protein-high versus protein-low cells by the matched RNA marker score. **(c-d)** Spatial expression maps of *EPCAM* (**c**) and *COL1A1* (**d**) **(e)** Per-gene Pearson correlation between predicted transcript assignments and Visium HD ground truth for MultiTME and iStar. **(f)** Per-pathway Pearson correlation between Hallmark gene set enrichment scores computed from MultiTME transcript assignments and from Visium HD, across 14 cancer-relevant pathways.
